# Island demographics and trait associations in white-tailed deer

**DOI:** 10.1101/2023.08.01.551454

**Authors:** Brooklyn S. Cars, Camille Kessler, Eric A. Hoffman, Steeve D. Côté, Daniel Koelsch, Aaron B.A Shafer

**Affiliations:** Environmental and Life Sciences Graduate Program, Trent University, 2140 East Bank Drive, Peterborough, ON K9J 7B8, Canada; Department of Forensics, Trent University, 2140 East Bank Drive, Peterborough, ON K9J 7B8, Canada; Department of Biology, University of Central Florida, 4000 Central Florida Blvd, Orlando, FL, USA; Département de Biologie and Centre d’Études Nordiques, Université Laval, G1V 0A6, Québec, QC, Canada; Fédération des chasseurs de Saint-Pierre et Miquelon. Saint-Pierre et Miquelon, France; Direction des Territoires de l’Alimentation et de la Mer, service Biodiversité. Saint-Pierre et Miquelon, France

## Abstract

When a population is isolated and composed of few individuals, genetic drift is the paramount evolutionary force that results in the loss of genetic diversity. Inbreeding might also occur, resulting in genomic regions that are identical by descent, manifesting as runs of homozygosity (ROHs) and the expression of recessive traits. Likewise, the genes underlying traits of interest can be revealed by comparing fixed SNPs and divergent haplotypes between affected and unaffected individuals. Populations of white-tailed deer (*Odocoileus virginianus*) on islands of Saint Pierre and Miquelon (SPM, France) have high incidences of leucism and malocclusions, both considered genetic defects; on the Florida Keys islands (USA) deer exhibit smaller body sizes, a polygenic trait. Here we aimed to reconstruct island demography and identify the genes associated with these traits in a pseudo case-control design. The two island populations showed reduced levels of genomic diversity and a build-up of deleterious mutations compared to mainland deer; there was also significant genome-wide divergence in Key deer. Key deer showed higher inbreeding levels, but not longer ROHs, consistent with long-term isolation. We identified multiple trait-related genes in ROHs including *LAMTOR2* which has links to pigmentation changes, and *NPVF* which is linked to craniofacial abnormalities. Our mixed approach of linking ROHs, fixed SNPs and haplotypes matched a high number (∼50) of a-priori body size candidate genes in Key deer. This suite of biomarkers and candidate genes should prove useful for population monitoring, noting all three phenotypes show patterns consistent with a complex trait and non-Mendelian inheritance.

## Introduction

Genomic architecture is the organized arrangement of genes and other functional elements underlying trait variation (Koonin 2009). Targeted population sampling combined with high-throughput sequencing can characterize the architecture and effect size of genomic variation underlying traits (Ceballos *et al*. 2018). Specifically, the genomes of individuals with and without traits of interest can be compared for the identification of divergent single nucleotide polymorphisms (SNPs) (Yang *et al*. 2008; Russell *et al*. 2012; Getmantseva *et al*. 2020). Reconstructing haplotypes is also useful for developing diagnostic tools and identifying candidate genes, with clear regions underlying body size variation in dogs detected for example (Parker *et al*. 2009). Large effect SNP and haplotypes, especially for polygenic traits, are typically found in low frequency (e.g. Yengo *et al*. 2022), and often require targeted phenotypic sampling to have enough power to detect in natural populations (e.g. Anderson *et al*. 2022).

The distribution of SNPs present in the genome is also useful for the identification of regions with reduced heterozygosity and segments of autozygosity (Kim *et al*. 2013); long rows of sequential homozygous sites known as runs of homozygosity (ROHs) can be used to map the genetic regions of recessive traits stemming from deleterious mutations (Hildebrandt *et al*. 2009). Runs of homozygosity result from the passing of identical haplotypes from one generation to the next as a result of mating between relatives, with the length of ROH corresponding to recency of the common ancestor (McQuillan *et al*. 2008; Kirin *et al*. 2010). Expression of recessive, often deleterious, traits due to homozygosity require ROHs to be present (Robinson *et al*. 2023), though purging might remove deleterious alleles (Wootton *et al*. 2023). A populations’ demographic history, notably recent trends in effective size (*N_e_*), also can correspond to ROH dynamics (Kardos *et al*. 2023) and estimates of ROH are now commonly included in assessments of endangered species, with for example unmanaged scimitar-horned oryx (*Oryx dammah*) populations manifesting significantly higher inbreeding (F_ROH_) than those in captivity (Humble *et al*. 2023). Therefore, detecting and mapping ROHs in a population can identify the genomic impact of demographic change and map genes that are associated with recessive and deleterious traits.

White-tailed deer (*Odocoileus virginianus*) are herbivorous, hooved ungulates that are native to the western hemisphere, primarily North America, but their range has expanded through human assistance to parts of Europe and various remote islands (Hewitt 2011; Villanova *et al*. 2017; Fuller *et al*. 2019; Entfellner *et al*. 2021). Anthropogenic movement appears to have impacted genetic load in white-tailed deer (Wootton *et al*. 2023), meaning the accumulation of deleterious alleles, but overall, population differentiation (e.g. Cullingham *et al. 2*011) and inbreeding across the species is low (F_IS_ <0.10; Hamlin *et al*. 2021). In some cases, introduced island populations show high diversity (Fuller *et al. 2*019; Nelson *et al. 2*021), but ensuing island dynamics have resulted in various phenotypic variants or anomalies in some deer populations. For example, the native population of white-tailed deer that inhabit the Florida Keys - referred to as Key deer - are significantly smaller compared to the white-tailed deer populations on the mainland (Maffei *et al*. 1988; Folk and Klimstra 1991; Karns 2010). Island populations of deer often experience reduced size and body mass from nutrition limitations (e.g. Anouk Simard *et al*. 2008; Bartareau 2019), but Key deer weigh less than the Florida mainland (Bartareau 2019) and are considered the smallest subpspecies (Hewitt 2011). These patterns are consistent with the Island rule of larger animals becoming smaller (Foster 1964; Benítez-López *et al*. 2021), but also follow a general clinal decline in deer body sizes at southern latitudes (Wolverton *et al*. 2009). Stature is generally a complex trait driven by many large and small effect genes (Kemper *et al*. 2012; Sidorenko *et al*. 2022) and Anderson *et al*.(2022) identified a suite of candidate genes impacting body size in deer (See Supplemental File S1). Key deer are genetically differentiated (Villanova *et al*. 2017), making disentangling drift from selection, and the Island rule, particularly challenging without first reconstructing their demographic history. Mark-recapture estimates place Key deer at 1,000 individuals (Villanova *et al. 2*017) and they are listed as Endangered under the Endangered Species Act.

In contrast, the introduced population of white-tailed deer on the islands of Saint Pierre and Miquelon (SPM) have relatively high rates of leucism and malocclusions, meaning incorrect alignment of the maxillary bone and the mandible. This population stems from 14 animals being introduced in the 1953; it is likely that the genes underlying these traits exhibit an increased frequency due to their presence in a small founding population, though inbreeding has not been assayed. While white-tailed deer are normally a uniform reddish-brown-grey with white fur scattered on appendages, face, and rear, deer with leucism present a partial to entire lack of pigmentation. Coat colour composition in ungulates is mainly determined by loci that control melanocyte activity and distribution (Fontanesi *et al*. 2010). Mutations in the melanocortin receptor locus (*MC1R*) in particular have been found to be major contributors to leucism (Haase *et al*. 2013; Reiner *et al*. 2020). Brachygnathism and prognathism are both forms of malocclusion (Faragalla 2000; Guan *et al*. 2015) and have been observed in both mule deer (*O. hemionus,* Robinette and Aldous 1955) and white-tailed deer (Smits and Bubenik 1990). Candidate genes have been identified for both leucism (Supplementary Table S1) and jaw malocclusions (Supplementary Table S2); leucism is considered a recessive trait (Pereira *et al*. 2023) but malocclusion inheritance patterns are less clear (Cakan *et al*. 2012). Expression of both traits in deer likely comes with a fitness cost (Wolff *et al*. 1993; Faragalla 2000; Reissmann and Ludwig 2013).

The emergence of divergent phenotypes (Key deer) and traits with a fitness cost (SPM) on islands present a pseudo case-control design allowing for characterizing the genetic architecture of phenotypes. By leveraging these phenotypically distinct populations and comparing to a large mainland white-tailed deer genomic data set (Kessler *et al*. 2023), we aim to reconstruct each population’s recent demographic history and genetic load, and test the prediction that ROH and candidate genes are associated with expression of recessive traits in SPM, while unique haplotypes with candidate body size genes underly the divergence of Key deer.

## Materials and Methods

### Sampling and Sequencing

Phenotype information for leucism or malocclusions from SPM was obtained from local managers and photos were examined to select individuals with one or more of the visible traits of interest (see Supplemental Fig. S1). Twenty-six white-tailed deer tissue samples were then selected from the two island populations of interest: SPM (France, but coastal Canada) and the Florida Keys (United States) in addition to four samples from Anticosti Island, Quebec (Canada). We leveraged available genome sequence data from four additional Key deer, one SPM sample (normal phenotype), and 21 deer from the North American mainland (normal phenotypes) that are part of a larger project (Fig. 1; PRJNA830519). Samples were provided by hunters voluntarily, with some collected from road mortality. DNA extraction from the tissue samples was carried out using the QIAGEN DNeasy Blood & Tissue Kit following the manufacturer’s instructions (*DNeasy® Blood & Tissue Handbook* 2020). We quantified the DNA concentrations using the Invitrogen Qubit assay. Samples were sent to The Centre for Applied Genomics in Toronto, Canada for low coverage (∼4x) whole genome sequencing (WGS) on an Illumina NovaSeq.

**Fig. 1:**
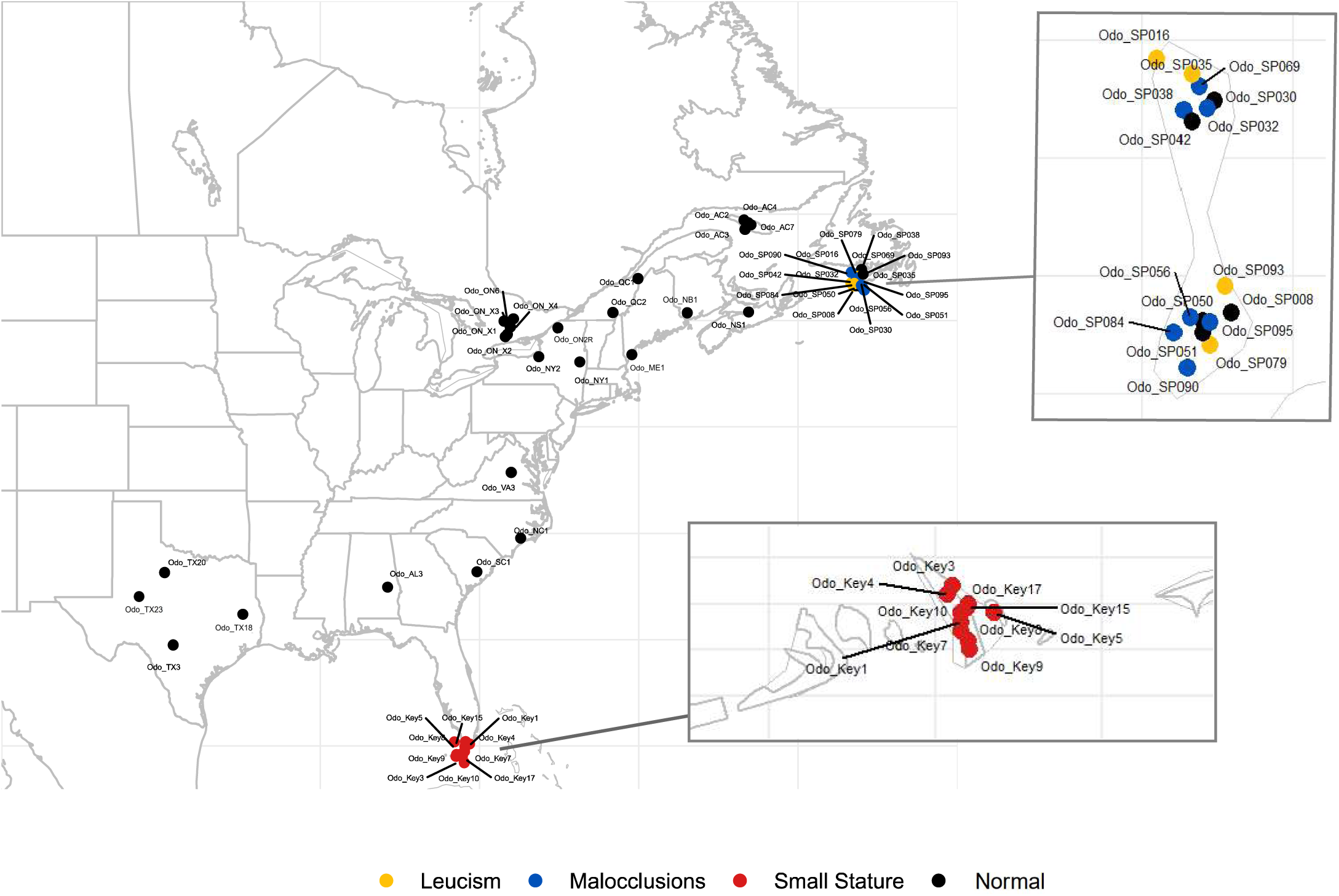
Sampling locations with phenotype represented by colours: Normal (black), Leucism (yellow), Malocclusion (blue), and Small Stature (red).

### Data Preparation

The quality of the raw data libraries was examined using FastQC (v0.11.9; Andrews 2010) and regions of poor quality were removed using Trimmomatic (v0.39; Bolger, Lohse and Usadel 2014). The trimmed sequence data was then re-examined using FastQC, mapped to the annotated *O. virginianus* reference genome (GCA_023699985.1; Anderson *et al*. 2022) using BWA-MEM (v0.7.17; Li 2013), and sorted using SAMtools sort (v1.15.1; Li *et al*. 2009). We detected duplicated reads using Picard MarkDuplicates (v2.23.2; Broad Institute, GitHub Repository 2019) and removed the duplicated reads using Sambamba (v0.8.2; Tarasov *et al*. 2015). Some samples had multiple sequence files and were merged before removing the duplicated reads using SAMtools merge. To ensure that no errors had been made throughout the process and that we had the desired coverage and depth to continue analysis we quantified metrics with SAMtools coverage and depth, and sambamba flagstat.

SNPs were called using ANGSD (v0.939; Korneliussen, Albrechtsen and Nielsen 2014) following the GATK model (-gl 2) to estimate genotype likelihoods (GLs) in a beagle genotype likelihood format (-doGlf 2). The genotypes were printed (-doGeno 4) and required SNPs with a minimum p-value of 1e−6 (-SNP_pval 1e-6), a minimum mapQ quality of 20 (-minMapQ 20), and a minimum base quality of 20 (-minQ 20). Further, we assumed that the major and minor alleles were known (-domaf 1) and that they can be inferred directly from likelihoods (- doMajorMinor 1).

### Estimating Subpopulation Diversity and Demography

We estimated the nucleotide diversity (π) among each subpopulation of our dataset using VCFtools (v0.1.16; Danecek *et al*. 2011). We calculated π using a 10 kb step size in between windows (-window-pi 10000). Tajima’s *D* for each subpopulation was in windows of 10 kb (- TajimaD 10000) using only bi-allelic sites. We estimated fixation indices (*F*_ST_) between populations using window sizes of 10 kb (-fst-window-size 10000). Using plink (v1.9) we generated a PCA based on the variance-standardized relationship matrix for the visualization of population stratification of the subpopulations. All plots were done in R version 4.2.2 (R Core Team 2022). To estimate the historical *N_e_*within the last 200 generations, we used the program GONE that uses the linkage disequilibrium information for markers across the genome (Santiago *et al*. 2020). We selected a maximum of 30,000 SNPs per chromosome, 0.5 cM/Mb (Kessler *et al*. 2023), an *hc* of 0.04 (maximum recombination rate), and applied the Haldane correction for genetic distance; we repeated the analysis 500 times. For a contemporary point estimate of *N_e_* we ran *currentNe* (Santiago *et al. 2*023) that extends the linkage disequilibrium approach but includes aspects of the species’ mating system. We ran *currentNe* with a random sample of 1,000,000 SNPs per island population with the mating structure parameter (*k*) inferred empirically from the data.

### Calling Runs of Homozygosity (ROHs)

Scaffolds making up the N90 were extracted using tabix (v.0.2.6.; Li 2011). We created tped and tfam files using plink (v2.0) and identified ROH genotypes using the plink window-based approach (v1.9). Data were lightly pruned for linkage disequilibrium (r^2^ > 0.90; Danecek *et al*. 2021). We used a sliding window size of 100 kb (-homozyg-kb 100), a minimum density of 1 SNP per 50 kb (-homozyg-density 50) and required a minimum of 50 SNPs to be considered a ROH (-homozyg-snp 50). We used the default scanning window size of 50 SNPs (-homozyg-window-snp 50), the default SNP hit rate of 0.05 (-homozyg-window-threshold 0.05) and allowed up to 3 heterozygote sites (-homozyg-window-het 3) as recommended for low coverage data (Ceballos *et al*. 2018) and 10 missing calls (-homozyg-window-missing 10) for each 100 kb window within called ROHs. No minor allele frequency cut-off was applied (Meyermans et al. 2020). A 1,000 bp gap between variant sites (-homozyg-gap 1000) was required for two ROHs to be considered separate and we grouped the samples in phenotypic pools to identify overlapping, and possibly matching ROH segments (-homozyg-group). The percentage of the genome covered by ROHs (F_ROH_) per individual was estimated by summing the total distance spanned by the ROH segments and dividing this by the total length of the N90 scaffolds.

We identified ROHs unique to each phenotypic group and found in >50% individuals, meaning none of the SPM normal, mainland or Anticosti Island comparators presented this ROH. Key deer we increased the required to >80% of samples have the ROH given the overall increased divergence of the population. The genomic coordinates of the ROHs were identified and matched to the *O. virginianus* gene annotation file (Anderson *et al*. 2022) and the biological function information of the discovered genes within ROH were identified through Uniprot (Pundir *et al*. 2016) and our candidate gene literature review (Supplementary Tables S1, S2, File S1). Any gene without an entry ID was screened on NCBI blast to identify a putative function. We inferred the approximate age of the identity by descent (IBD) event (no. of generations or *g*) based on ROH length, where *g* = 100/(2*r*L), with *r* being the recombination rate of 0.50 cM/Mbp (Kessler *et al*. 2023) length (L) of ROH in Mbp (Browning and Browning 2015; Kardos *et al*. 2017). To quantify whether our ROH-phenotype groupings could have occurred by chance, we permutated mainland, Anticosti and SPM individuals IDs and re-ran the analysis for leucism groupings 100 times.

### Fixed SNPs and haplotypes

We identified SNPs that were fixed in the phenotypic group, with the alternate allele found in >50% of wild-type samples (i.e. south mainland vs Key deer, SPM normal vs. leucism and malocclusions; Fig. 1). Using VCFtools (v0.1.16) we extracted lists of SNPs fixed in the local phenotypic group, requiring the alternate allele to be fixed in the mainland comparator. For SPM and the Key deer, we extracted and assayed all fixed SNPs. We complemented this with a formal haplotype analysis; here samples were first phased using SHAPEIT2 (Delaneau *et al*. 2013) then analyzed through a likelihood ratio test HapLRT (Garrison *et al*. 2022) to model the haplotype length distributions of the target (island) and background (comparator) populations; note the entire island sample of SPM was used to increase power. Here a positive sign from HapLRT reflect longer haplotypes in the target population; haplotype blocks were then defined as any region having a positive sign with successive p values <0.01 spanning 100 kb or longer and separated by at least 5 kb from other blocks. All fixed SNPs and haplotype blocks were matched to the *O. virginianus* gene annotation file (Anderson *et al*. 2022) and we identified genes within haplotype blocks or within 50 kb of the fixed SNP grouping. Lastly, for all phenotypic and mainland groupings we used SNPeff (Cingolani *et al*. 2012) to estimate and summarize the putative impact of all present SNPs; we focused on genetic load by measuring homozygote missense and loss of function (LOF) mutation, the latter being SNPs that resulted in a premature stop codon.

## Results

### Sampling, Sequencing and Data Preparation

All deer samples were sequenced to ∼4x coverage; this included 16 individuals from SPM of which four presented with leucism and seven with malocclusions, as well as 10 Key deer samples (smaller stature size). As comparators, we used 21 samples from the North American mainland split into North and South groupings and four from Anticosti Island, all of which were normal phenotypes (Fig. 1). We identified a total of 100,544,312 SNPs in all individuals and the analyses were performed exclusively on the N90 scaffolds that harboured 90,993,235 SNPs.

### Subpopulation Diversity

Tajima’s *D* for SPM was the highest followed by Key deer that had the lowest value of π (Table 1). Mainland and Anticosti deer showed negative Tajima’s *D* values. The fixation index (*F*_ST_) and PCA showed high levels of genetic differentiation between the Key deer and every other subpopulation; SPM formed its own cluster, while Anticosti Island was embedded within the North Mainland samples (Supplementary Table S3; Supplementary Fig. S2). Recent demographic trends revealed a stable long-term low *N_e_* of Key deer, with a large crash in SPM that predates the translocation (Fig. 2). Point estimates of current *N_e_*are low for both populations: *N_e_* of SPM (108; 90% confidence interval [CI] 64 –180) and Key deer (287; 90% CI 95–870).

**Fig. 2:**
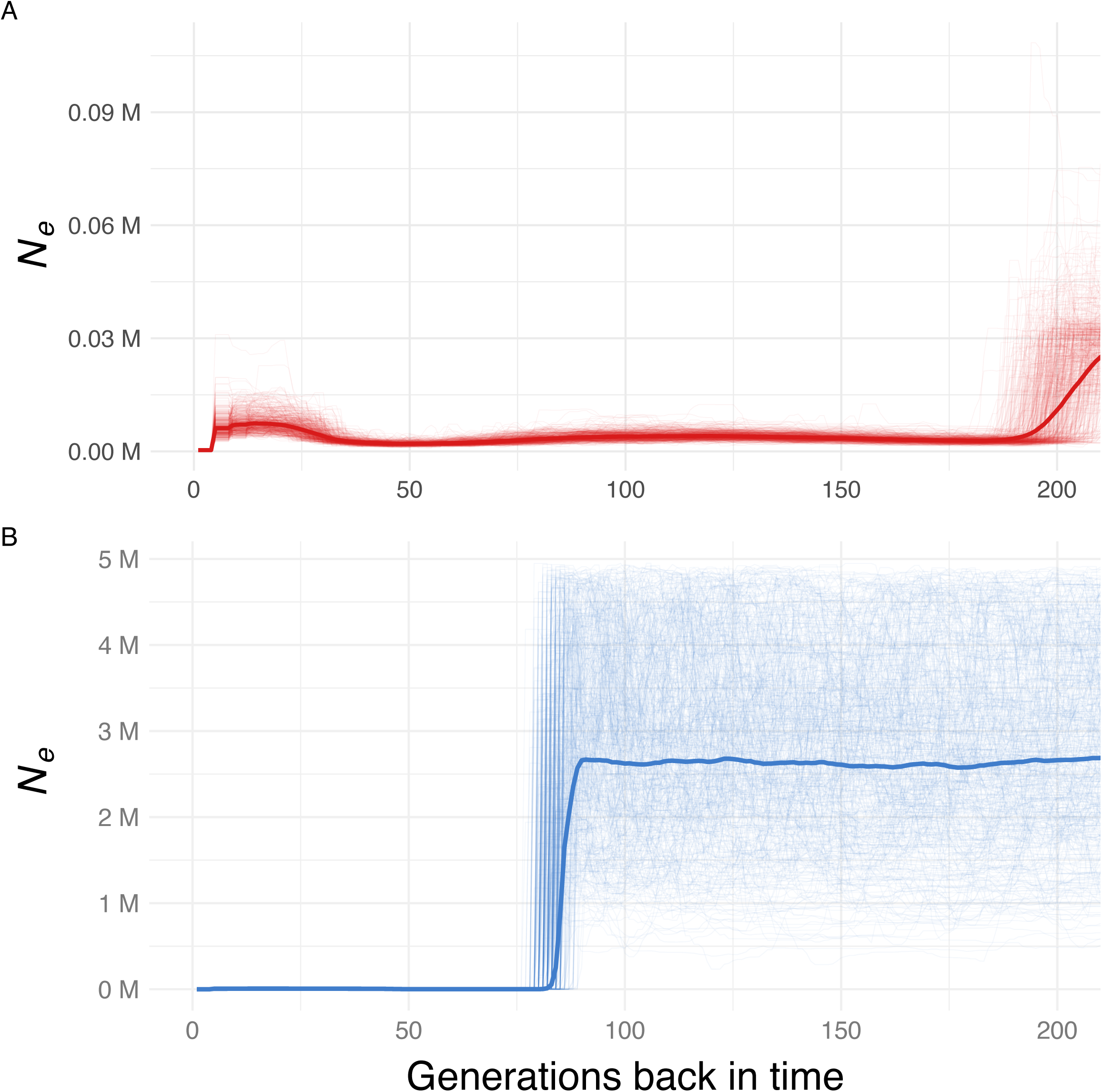
Effective population size changes for the last 400 years, x-axis as generations before present with a generation time of two years A) Key Deer B) Saint Pierre et Miquelon

**Table 1:** Summary statistics of subpopulations analyzed.

| Subpopulation | Tajima's $D$ | Nucleotide Diversity ($\pi$) | Mean $F_{\text{ROH}}$ | Range of average ROH length (kb) |
| --- | --- | --- | --- | --- |
| Anticosti | -0.09 | 0.007 | 0.01 | 153.41 – 172.89 |
| SPM | 0.54 | 0.005 | 0.02 | 150.92 – 202.66 |
| North Mainland | -0.39 | 0.006 | 0.01 | 151.85 – 233.50 |
| South Mainland | -0.51 | 0.007 | 0.01 | 158.69 – 191.03 |
| Florida Keys | 0.22 | 0.003 | 0.05 | 136.80 – 192.08 |

### Distribution of ROH among phenotypes

The populations with the highest F_ROH_ were the Key deer followed bhy SPM, while the populations from the North American Mainland and Anticosti Island all had a relatively low F_ROH_ (Fig. 3A). The range of average ROH lengths within the Key deer individuals was shorter than the other populations with ROH lengths of 136.8 kb – 192.1 kb (Table 1); this range of ROH likely corresponds to >500 generations to the most recent common ancestor. Additionally, all subpopulations had majority of their ROHs in the 100-300 kb size range (Fig. 3B), which corresponds to the greater total ROH length of this size range as well (Fig. 3C).

**Fig. 3:**
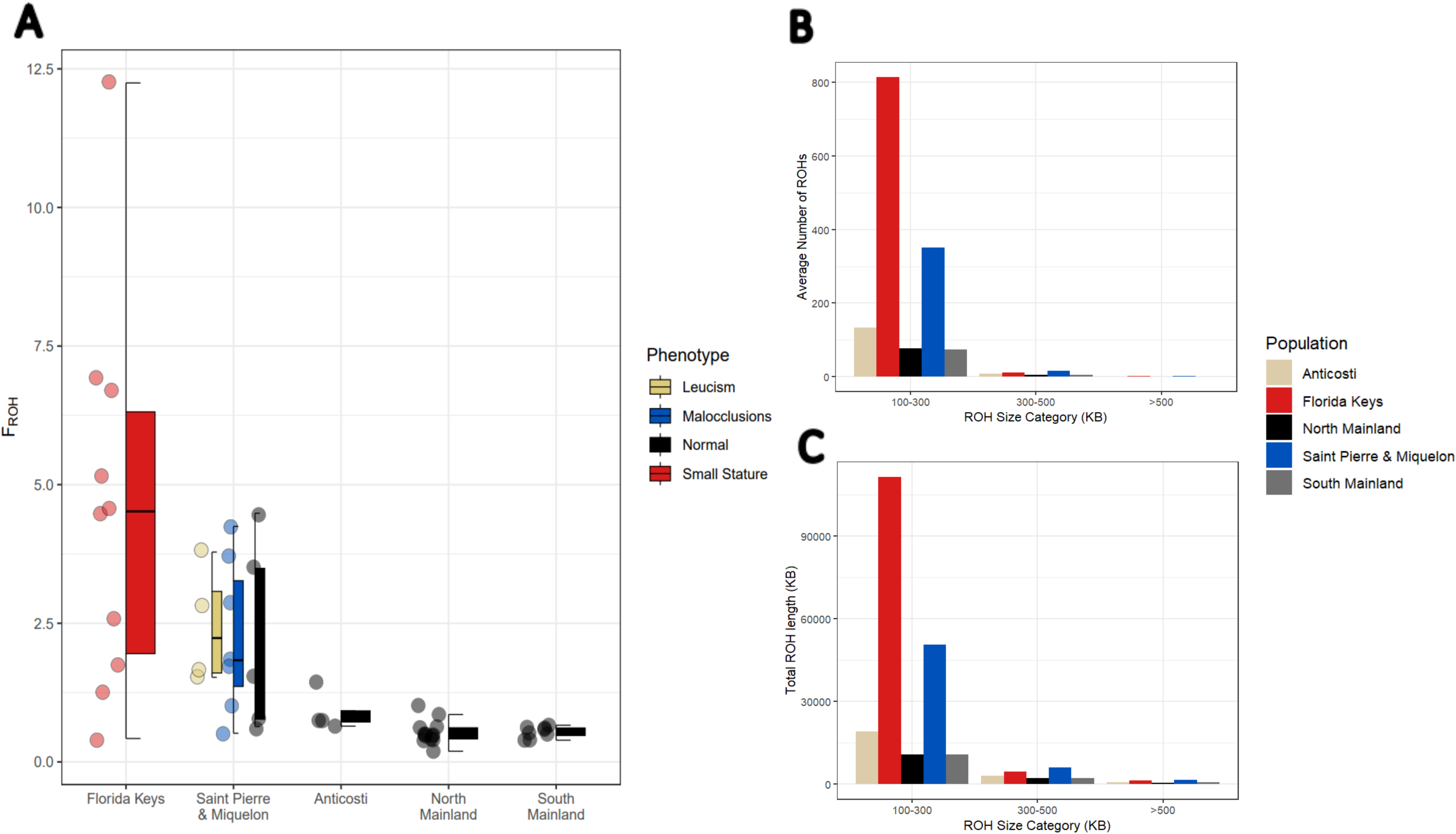
A) Percentage of the genome covered by ROHs (F_ROH_) for each population. B) Bar plot showing the total number of ROHs in each population stratified by ROH size category in kb. C) Bar plot showing the total length (kb) of all ROHs in each size category (kb).

In the pools for all four SPM white-tailed deer with leucism, we found 39 shared ROHs. Of those, one was unique to the individuals with leucism, and it contained 19 genes (Table 2); no a-priori candidate genes for leucism were detected (Supplementary Table S1). ROHs detected in animals with leucism and absent from the comparators contained multiple genes, but notably *LAMTOR2* (Endosomal adaptor protein p14) (Table 2). A total of 21 ROHs were found to be shared between all SPM white-tailed deer with malocclusions, but none were unique to this phenotype. One ROH was unique to 4/7 deer with this phenotype, and only harbored one gene (*NPVF*; Table 2). Lastly, 18 ROHs were shared among the smaller-stature Key deer. Of those, one was unique to the Key deer, and it contained 1 gene (*CHMP3*); which was also found within the Key deer’s fixed SNPs (Table 2; Supplementary File S2). Within a pool of all 10 Key deer and one SPM deer, there were two genes that matched the Anderson *et al*. (2022; Supplementary File S1) candidate gene table: *ART1* and *ART5* (Table 2). In addition, nine out of 10 of the Key deer shared eight ROHs that were unique to them, they collectively harboured 27 genes and had one gene that matched the Anderson *et al*.(2022) candidate gene table: *RPL17* (Table 2). Pools were unlikely to have arisen by chance (Table S4).

**Table 2:**
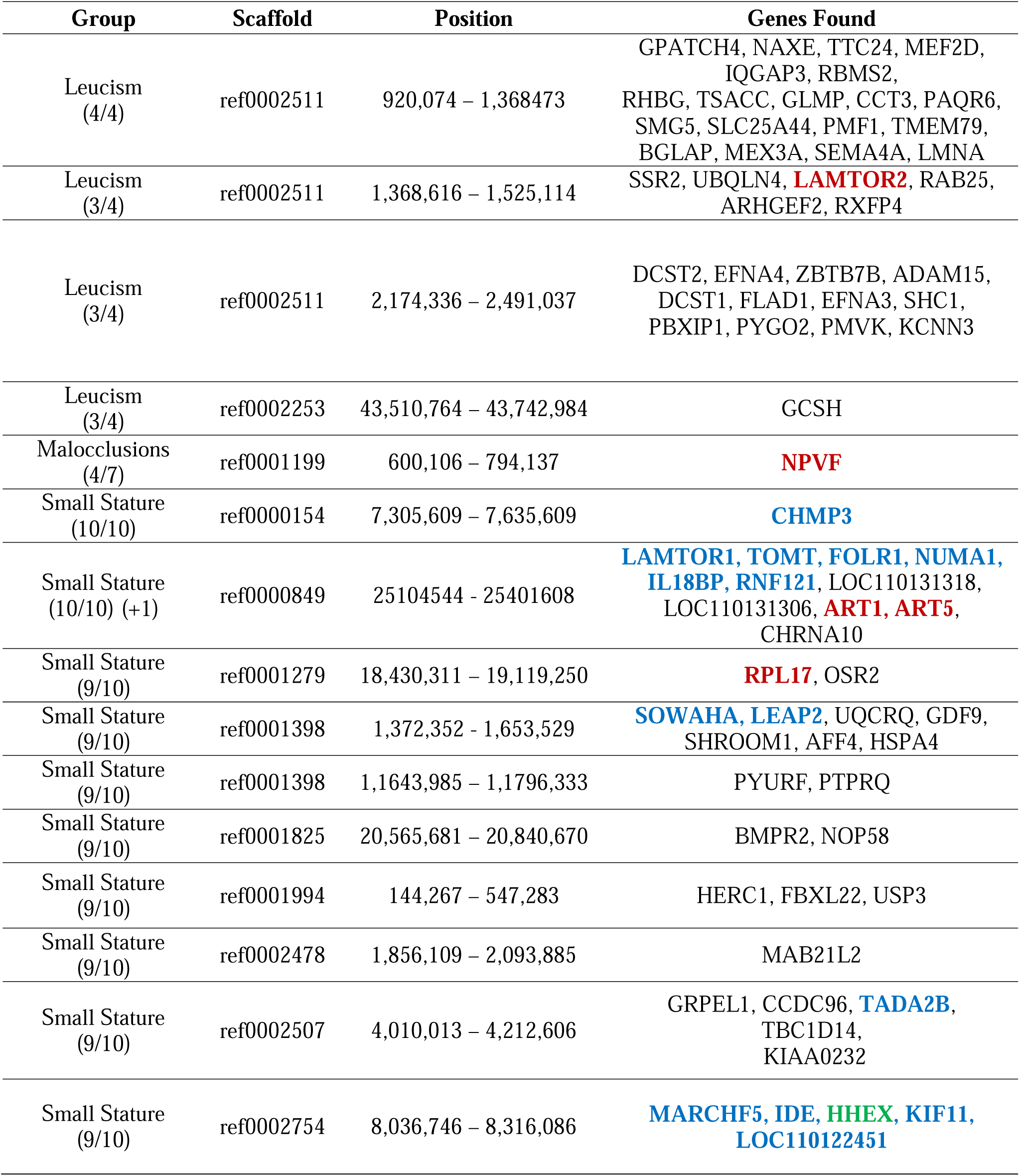
List of ROHs shared by subsets of each group and the genes found within them (Matching candidate gene table = red, fixed SNPs = blue, haplotypes = orange, and haplotypes and fixed SNPs = green)

### Fixed SNPs and haplotypes

For each fixed SNP we screened 50 kb flanking regions on either end to identify genes that were in close proximity; here we identified 72,291 fixed SNPs for Key deer population that collectively encompassed 9,081 genes. Of these, 20 overlapped with the ROHs and 427 matched the Anderson *et al*. (2022) candidate gene table. There were 159 fixed SNPs within the SPM leucism group that contained 65 genes, five fixed SNPs within the SPM malocclusion group that contained two genes and four fixed SNPs within the Anticosti population that contained one gene. There was no overlap for fixed SNPs and ROHs or candidate genes in either SPM phenotypic group. Fixed SNPs and ROH overlapped SNPs detected with a standard genome-wide association approach (Figure S4).

There were 695 haplotype blocks detected in Key deer, which encompassed 455 genes. One haplotype overlapped with the ROHs for Key deer, 384 overlapped with the fixed SNPs for Key deer, and 50 genes matched the candidate gene table, and notably included the gene *IGF1R* (Supplementary File S2). For the SPM population, we identified four haplotype blocks; these blocks harboured two genes, neither of which overlapped with the ROHs in SPM deer. The absolute distribution of variants, including deleterious, were similar amongst populations (Table S5). Homozygote LOF and missense mutations were notably elevated in Key deer (Fig. 4); there was a modest increase in SPM, specifically relative to Anticosti and mainland deer (Fig. 4).

**Fig 4:**
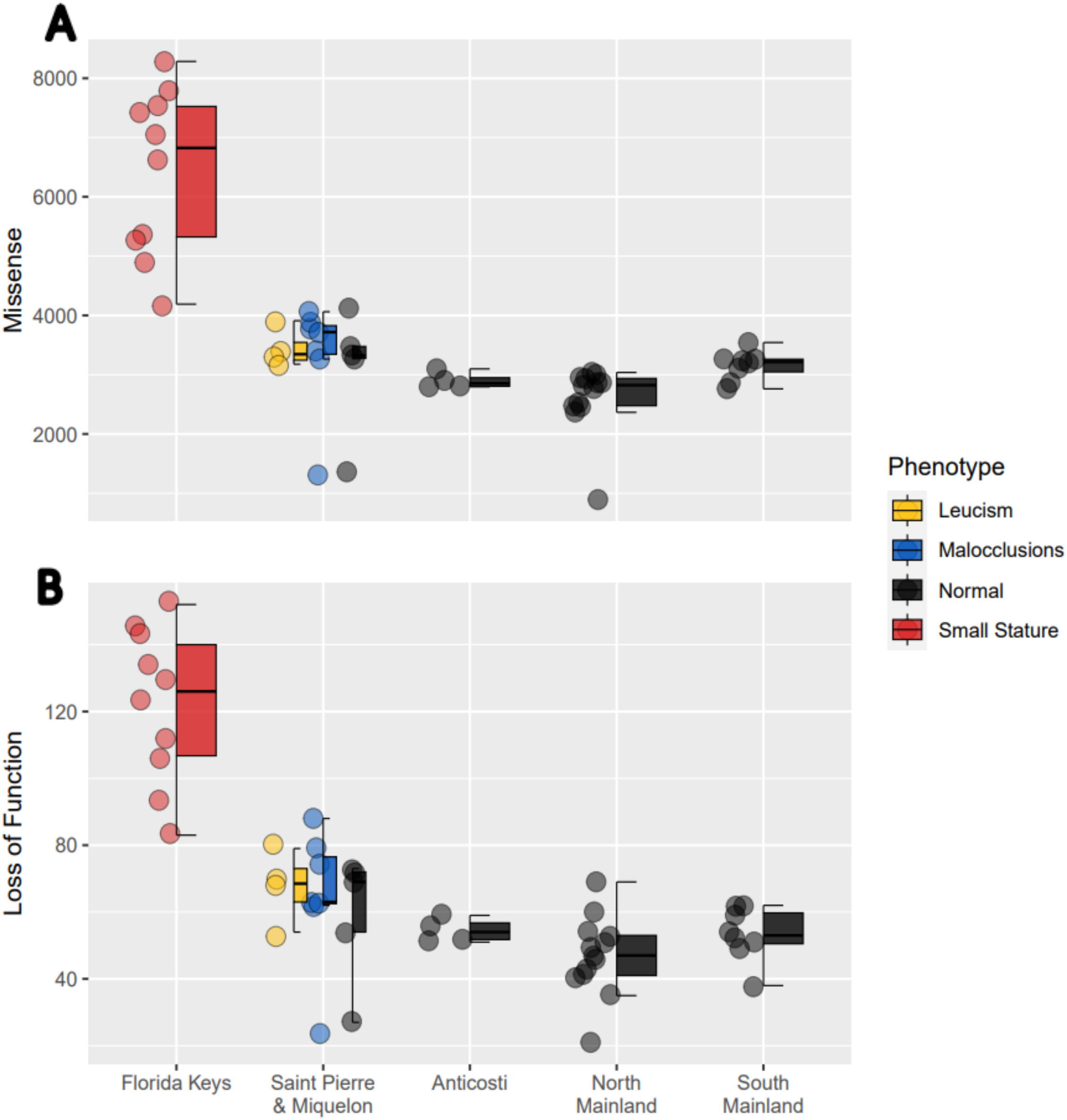
A) Number of homozygous missense mutations based on SNPeff annotations. B) Number of homozygous loss of function mutations based on SNPeff annotations. Populations are colour coded by phenotype: Normal (black), Leucism (yellow), Malocclusion (blue), and Small Stature (red).

## Discussion

Island populations are prone to drift; further, isolated populations composed of a small number of individuals are more likely to suffer from inbreeding, accumulate deleterious alleles, and manifest undesirable traits (Soule 1973; Villanova *et al*. 2017). Small populations, however, might not show inbreeding depression (e.g. Robinson *et al*. 2018), with purging occurring under the right demographic scenarios (Wootton *et al. 2*023). The presence of presumed deleterious phenotypes and the corresponding genetic data (e.g. Fig. 4), suggest that purging has not occurred in our focal populations. SPM deer have a relatively high frequency of malocclusions and leucism, and Key deer are significantly smaller than their mainland counterparts (Folk and Klimstra 1991). SPM samples clustered adjacent to northern mainland deer due to their recent translocation, while the Key deer are significantly diverged (see Supplementary Fig. S2). The Key deer became isolated approximately 8,000 years ago (Lazell 1989) while SPM were introduced in 1953; consequently, the confounding effects of drift are much less of a factor in the gene-trait associations detected in SPM compared to the Key deer. Still, clear patterns emerged that not only impact our understanding of the genetic architecture of traits in wild populations, but also the build-up of ROH and genetic load in light of recent demographic change.

### A tale of two island populations

Key deer are an endangered species under the US Federal Endangered Species Act. Genetic mark recapture studies suggests a population of ∼1,000 individuals, with an *N_e_* of only 11 (Villanova *et al*. 2017). Our genome wide estimates are higher, with a contemporary point estimate of ∼300. Here we suspect the cross-species amplification and small number of microsatellites resulted in a downward bias estimate in Villanova *et al*. (2017) compared to genome wide estimates based on linkage disequilibrium. Values of π, which are reflective of long-term or coalescent *N_e_* (Peart *et al. 2*020) are lowest in Key deer compared to all comparators; the coalescent *N_e_* / *N_C_*ratio does appear to be linked to conservation status and extinction risk (Peart *et al*. 2020; Dedato *et al*. 2022; Wilder *et al*. 2023); solving for π = 4*N_e_µ* and with *N_C_* estimates of Villanova *et al*.(2017) and *µ* of Kessler *et al*.(2023), would generate a ratio coalescent *N_e_*/ *N_C_* ratio >1 in Key deer, thereby supporting the endangered species designation.

Key deer were almost hunted to extinction in the 1950s (U.S. Fish and Wildlife Service 1999); despite this, the *N_e_* has been stable for >200 generations (Fig. 2A) and no signal of overharvest is detected, as seen in mainland deer (Kessler and Shafer 2023). Collectively, this suggests the moderate buildup of ROH (Fig. 3A), and genetic load (Fig. 4) is due to the historical demographic bottleneck, and not to contemporary inbreeding or overharvest. Indeed, while Key deer have the highest mean F_ROH_ of white-tailed deer examined (Table 1), the average length of ROH is not longer, with the number of generations to the common ancestor of the ROH being >500. And while there is an elevated genetic load, with values higher than relevant comparators (Humble *et al*. 2023), there does not appear to be a link to population viability given the general patterns seen in deer (Kessler and Shafer 2023) and robust population growth of white-tailed deer in general (Webb 2018). The relatively high *N*_e_ in Key deer might also have limited purging, as seen in ungulates with much lower *N*_e_ (e.g. mountain goats; Wootton *et al. 2*023; Martchenko and Shafer 2023). Scandinavian wolves, for example, show most ROH due to recent ancestry but also high variance in estimates with F_ROH_ ranging from 0.01 to 0.54 (Kardos *et al. 2*018). Accordingly, deleterious alleles have also become fixed (Smeds and Ellegren 2023). Here the demographic collapse in wolves was considerably worse than Key deer, and recovery (or growth) much slower which would explain these differences. Collectively, these data support Villanova *et al*. (2017) who predicted long-term persistence of the Key deer population.

The SPM Island population originated from 14 white-tailed deer; in contrast 220 white-tailed deer were introduced to the much larger Anticosti Island (7,953 km²) off the coast of Quebec in 1896 (Fuller *et al*. 2019). Accordingly, diversity values (Table 1) reflect this difference in founding number and island size, with Anticosti Island largely indistinguishable from the mainland. Tajima’s *D* values were also highest on SPM, suggesting a recent loss of rare alleles due to this founding effect. The crash in recent *N_e_*timepoints (Fig. 2B) predates the translocation event; as genomes carry a historical record of the species, this signal is a carry-over from the source population and colonial overharvest on the mainland (Kessler and Shafer 2023). There does, however, appear to be a slight build-up of genetic load from the translocation event (Fig. 4). In island simulations, Robinson *et al*. (2018) showed that strongly deleterious alleles were purged, while moderate deleterious alleles accumulated. It is therefore conceivable that the genetic load seen in both Key deer and SPM consists primarily of moderate to low effect deleterious mutations allowing their persistence; temporal monitoring of SPM will help shed light on this important evolutionary process given the recency of the founding event.

### Inbreeding and isolation lead to detecting candidate genes

The Islands of SPM have a relatively high number of individuals with leucism and malocclusions. None of the candidate genes for leucism (Hauswirth *et al*. 2012; Haase *et al*. 2013; Reiner *et al*. 2020) were found within our ROH segments. There were 19 new candidate genes harboured by the ROHs unique to all leucism individuals, though none had a clear biological link. Screening ROHs shared by 3/4 animals in the leucism pool identified the gene *LAMTOR2* (Endosomal adaptor protein p14) which has links to hypopigmentation (Bohn *et al*. 2007). Hypopigmentation is caused by the hindrance of melanocytes which are responsible for the pigmentation of hair and skin (Dessinioti *et al*. 2009). *LAMTOR2* interacts with the *MITF* transcription factor which regulates melanocyte activity and development; as leucism is known to be caused by mutations in genes that are associated with melanocytes *MITF* could be the leucism pathway in white-tailed deer. Our results also hint at non-Mendelian patterns of inheritance as suggested in other vertebrates (Edelaar *et al*. 2011; Jablonski *et al*. 2017). No ROHs were found to be unique to all of the white-tailed deer with malocclusions; however, a unique ROH segment that was shared by 4/7 white-tailed deer with malocclusions contained the gene *NPVF*. In humans and mice enhancers adjacent to the *NPVF* are implicated in craniofacial abnormalities (Wildermann *et al*. 2022; Yu *et al*. 2023). While this region certainly appears functionally relevant, not all animals with maloclussions have this ROH, suggesting again a polygenic trait or incomplete penetrance (Stiles and Luke 1953). Kardos *et al*. (2018) suggested that areas with low ROH abundance are likely to contain loci with deleterious recessive alleles that contribute disproportionately to inbreeding depression. Thus, we cannot rule out, homozygous deleterious variants that are not captured by our ROH quantification.

Stature size is a complex trait influenced by many potential genes that have both large and small effects. Anderson *et al*. (2022) sequenced the genomes of extreme phenotypes in deer, specifically the tail ends of the distribution for body size controlling for age and created an extensive list of candidate genes (Supplemental File S1). Our study found thousands of fixed SNPs, clearly due to drift and long-term isolation. Still, close to 50 genes that overlapped previous genome-association work (Anderson *et al. 2*022) fell on both haplotypes and fixed SNPs suggesting these are indeed likely linked to body size. Body size is clearly polygenic and it would be tempting to create a story of how these genes influence body size (Pavlidis *et al*. 2012), so we simply note that one haplotype block contained the gene *IGF1R* which has been seen to cause a reduced size in dogs (*Canis familiaris*, Hoopes *et al*. 2012), which in our system is consistent with the island rule.

## Conclusion

Island white-tailed deer populations have arisen from a mix of human-assisted and natural colonization or isolation. Demographic assessment shows a stable, but low *Ne* in both SPM and Key deer, an accumulation of genetic load, but generally low levels of ROH. Our results could be useful for wildlife management concerned with small and isolated populations that may be experiencing a loss of genetic diversity due to inbreeding. Notably, combining Tajima’s *D* with demographic models, and ROH with genetic load metrics, helps elucidate the role of demographic change in shaping population genomic health. Interestingly, the island populations do not have longer ROH; thus, expression of putative recessive traits on SPM is not due to recent inbreeding, rather chance autozygosity due to a high frequency of the gene variant(s) in the founding population. Collectively, this supports more detailed demographic models of Key suggesting they have been isolated for a considerable length of time (Kessler *et al*. 2023), with this study showing no evidence of recent inbreeding, despite an increased genetic load.

Our mixed approach of ROH, fixed SNPs and haplotypes did pull out a-priori candidate genes and discovered others. Storytelling of gene ontologies is common and leads to false positives (Pavlidis *et al*. 2012), but our findings support all three traits not having a simple architecture, and identified promising genes involved with leucism (*LAMTOR2*), malocclusions (*NPVF*), and stature size (*IGF1R*) in white-tailed deer. Despite the divergence in Key deer limiting some approaches (i.e. fixed SNPs were associated with almost half the total genes suggesting a large impact of genetic drift), both haplotype and ROH inferences overlapped with a large number of candidate genes allowing us to weed out background noise. Given the relatively small sample sizes, the validation of autozygosity segments and quantification of effect sizes in independent samples and mouse models will be required to validate these genes.

## Acknowledgements

Participation from hunters, managers and community members is essential to this research. We want to thank the community of St-Pierre and Miquelon that provided samples. Additional Florida Key samples were provided by George Zaragoza; the remaining providers are found listed in (Kessler *et al*. 2023). BC was supported by a Natural Sciences and Engineering Research Council (NSERC) Undergraduate Student Research Award and the Trent Summer Work Experience Program. This work was supported by NSERC Discovery Grant (ABAS; SDC); ComputeCanada Resources for Research Groups (ABAS); Canadian Foundation for Innovation (ABAS): John R. Evans Leaders Fund (ABAS) and the Ontario Early Researcher Award (ABAS). We thank four anonymous referees and the Editor for valuable comments. Trent University is located on the traditional territory of the Michi Saagiig Anishnaabeg.

## Author contribution statement

BC and ABAS conceived the study, BC, CK, EH, SDC, DK, ABAS collected the samples, BC and CK performed the molecular laboratory work and the bioinformatic analyses with contribution from ABAS. BC and ABAS wrote the initial draft of the manuscript, all authors reviewed and commented on the manuscript.

## Data archiving

Raw reads for the 25 deer additional individuals were deposited on the NCBI Sequence Read Archive (Accession number PRJNA830519). All scripts are available on GitLab (https://gitlab.com/WiDGeT_TrentU/undergrad-theses/-/tree/master/%20Cars_2023)

## Ethics approval

All experiments have been conducted as per the guidelines of the Trent University Research Ethics Board. No approval of research ethics committees was required to accomplish the goals of this study because sequencing work was conducted on samples donated by hunters or collected as roadkill.

