## Supplemental Tables and Figures for "Island demographics and trait associations in white-tailed deer"

**Table S1:** Leucism candidate genes

| Gene Name | References |
|---|---|
| TYR | (Reiner, Tramberend, *et al.*, 2020) |
| MITF | (Haase *et al.*, 2013; Hofstetter *et al.*, 2019) |
| PAX3 | (Hauswirth *et al.*, 2012) |
| KIT | (Haase *et al.*, 2013) |
| EDNRB | (Wang *et al.*, 2015) |
| ASIP | (Xu *et al.*, 2015) |
| BRM | (Xu *et al.*, 2015) |
| MC1R | (Reiner, Weber, *et al.*, 2020) |
| PMEL | (Liu *et al.*, 2022) |
| LAMTOR2 | (Bohn *et al.*, 2007) |
| LVRN (Taqpep) | (Kaelin *et al.*, 2012) |

**Table S2:** Malocclusion candidate genes

| Gene Name | References |
|---|---|
| MATN1 | (Rodrigues *et al.*, 2013) |
| DUSP6 | (Nikopensius *et al.*, 2013) |
| FGF23 | (Chen *et al.*, 2015) |
| ADAMTS1 | (Guan *et al.*, 2015) |
| NPVF | (Wildermann *et al.*, 2022; Yu *et al.*, 2023). |
| ARHGAP21 | (Perillo *et al.*, 2015) |
| MYO1H | (Sun *et al.*, 2018) |
| IGF1 | (Xue, Rabie and Luo, 2014) |
| COL2A1 | (Xue, Rabie and Luo, 2014) |

**Table S3:** $F_{ST}$ calculated between each subpopulation of white-tailed deer (0.05 – 0.15 = moderate genetic difference, 0.15 – 0.25 great genetic difference, >0.25 = very great genetic difference)

| | **Anticosti** | **SPM** | **North Mainland** | **South Mainland** |
|---|---|---|---|---|
| **St-Pierre and Miquelon** | 0.07 | | | |
| **North Mainland** | 0.01 | 0.06 | | |
| **South Mainland** | 0.04 | 0.10 | 0.02 | |
| **Florida Keys** | 0.38 | 0.35 | 0.29 | 0.30 |

**Table S4:** Permutation test of leucism individuals (100% denotes ROHs shared between all leucism individuals, 75% is ¾ leucism individuals etc.).

| File # | 100% | 75% | 50% |
|---|---|---|---|
| Original | 1 | 3 | 35 |
| 1 | 0 | 0 | 0 |
| 2 | 0 | 0 | 1 |
| 3 | 0 | 0 | 2 |
| 4 | 0 | 0 | 2 |
| 5 | 0 | 0 | 1 |
| 6 | 0 | 0 | 6 |
| 7 | 0 | 0 | 6 |
| 8 | 0 | 0 | 2 |
| 9 | 0 | 0 | 7 |
| 10 | 0 | 0 | 3 |
| 11 | 0 | 0 | 3 |
| 12 | 0 | 0 | 10 |
| 13 | 0 | 0 | 9 |
| 14 | 0 | 1 | 4 |
| 15 | 0 | 0 | 0 |
| 16 | 0 | 0 | 0 |
| 17 | 0 | 1 | 3 |
| 18 | 0 | 0 | 15 |
| 19 | 0 | 0 | 2 |
| 20 | 0 | 0 | 2 |
| 21 | 0 | 0 | 3 |
| 22 | 0 | 0 | 12 |
| 23 | 0 | 0 | 2 |
| 24 | 0 | 0 | 1 |
| 25 | 0 | 0 | 15 |
| 26 | 0 | 0 | 0 |
| 27 | 0 | 0 | 1 |

| File # | 100% | 75% | 50% |
|---|---|---|---|
| 58 | 0 | 0 | 1 |
| 59 | 0 | 0 | 0 |
| 60 | 0 | 0 | 0 |
| 61 | 0 | 0 | 0 |
| 62 | 0 | 0 | 4 |
| 63 | 0 | 0 | 3 |
| 64 | 0 | 0 | 2 |
| 65 | 0 | 0 | 0 |
| 66 | 0 | 0 | 1 |
| 67 | 0 | 0 | 1 |
| 68 | 0 | 0 | 22 |
| 69 | 0 | 0 | 0 |
| 70 | 0 | 0 | 0 |
| 71 | 0 | 0 | 14 |
| 72 | 0 | 0 | 3 |
| 73 | 0 | 0 | 1 |
| 74 | 0 | 0 | 1 |
| 75 | 0 | 0 | 8 |
| 76 | 0 | 0 | 14 |
| 77 | 0 | 0 | 0 |
| 78 | 0 | 0 | 3 |
| 79 | 0 | 1 | 16 |
| 80 | 0 | 0 | 4 |
| 81 | 0 | 1 | 3 |
| 82 | 0 | 1 | 41 |
| 83 | 0 | 0 | 2 |
| 84 | 0 | 0 | 5 |
| 85 | 0 | 0 | 3 |

| | | | |
|---|---|---|---|
| 28 | 0 | 0 | 3 |
| 29 | 0 | 0 | 2 |
| 30 | 0 | 0 | 2 |
| 31 | 0 | 0 | 0 |
| 32 | 0 | 0 | 17 |
| 33 | 0 | 0 | 1 |
| 34 | 0 | 0 | 2 |
| 35 | 0 | 0 | 1 |
| 36 | 0 | 0 | 4 |
| 37 | 0 | 0 | 25 |
| 38 | 0 | 1 | 25 |
| 39 | 0 | 0 | 1 |
| 40 | 0 | 2 | 11 |
| 41 | 0 | 1 | 35 |
| 42 | 0 | 0 | 6 |
| 43 | 0 | 0 | 1 |
| 44 | 0 | 0 | 1 |
| 45 | 0 | 1 | 18 |
| 46 | 0 | 0 | 2 |
| 47 | 0 | 0 | 0 |
| 48 | 0 | 0 | 2 |
| 49 | 0 | 1 | 1 |
| 50 | 0 | 0 | 1 |
| 51 | 0 | 2 | 6 |
| 52 | 0 | 0 | 0 |
| 53 | 0 | 0 | 0 |
| 54 | 0 | 0 | 1 |
| 55 | 0 | 0 | 11 |
| 56 | 0 | 0 | 0 |
| 57 | 0 | 0 | 3 |

11

| | | | |
|---|---|---|---|
| 86 | 0 | 0 | 0 |
| 87 | 0 | 0 | 12 |
| 88 | 0 | 0 | 0 |
| 89 | 0 | 0 | 8 |
| 90 | 0 | 0 | 0 |
| 91 | 0 | 0 | 0 |
| 92 | 0 | 1 | 3 |
| 93 | 0 | 0 | 2 |
| 94 | 0 | 0 | 7 |
| 95 | 0 | 0 | 2 |
| 96 | 0 | 0 | 5 |
| 97 | 0 | 0 | 7 |
| 98 | 0 | 0 | 0 |
| 99 | 0 | 0 | 2 |
| 100 | 0 | 0 | 6 |

**Table S5:** snpEff Number of Effects by Impact and Functional Class. Numbers are relative to the variants assessed.

| Population | High | Low | Moderate | Missense | Nonsense | Silent |
|---|---|---|---|---|---|---|
| **Anticosti** | 2,522 (0.01%) | 141,028 (0.26%) | 56,143 (0.10%) | 56,572 (38.54%) | 1,216 (0.83%) | 88,997 (60.63%) |
| **St-Pierre and Miquelon** | 3,168 (0.01%) | 163,333 (0.25%) | 69,681 (0.11%) | 70,242 (39.56%) | 1,556 (0.88%) | 105,773 (59.57%) |
| **North Mainland** | 4,666 (0.01%) | 232,320 (0.26%) | 104,867 (0.12%) | 105,671 (39.47%) | 2,364 (0.88%) | 159,680 (59.65%) |
| **South Mainland** | 4,001 (0.01%) | 207,103 (0.26%) | 91,466 (0.11%) | 92,200 (38.86%) | 1,967 (0.83%) | 143,090 (60.31%) |
| **Florida Keys** | 2,437 (0.01%) | 88,465 (0.27%) | 44,145 (0.13%) | 44,473 (41.87%) | 1,360 (1.28%) | 60,386 (56.85%) |

**Fig. S1:** Local manager photos of leucism and malocclusion affected deer.

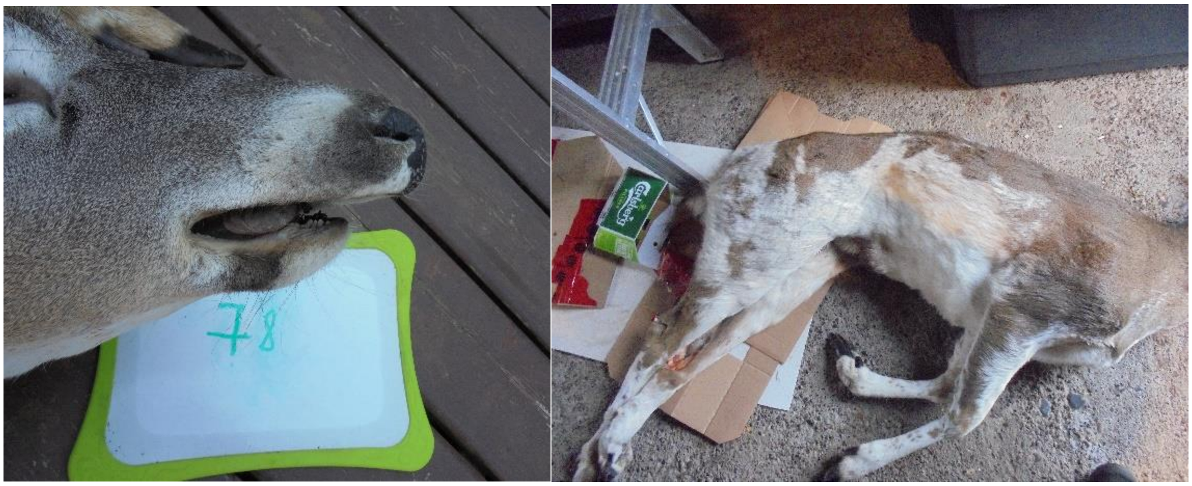

**Fig. S2:** Principal component analysis of subpopulation genetic variation. Populations are colour coded by phenotype: Normal (black), Leucism (yellow), Malocclusion (blue), and Small Stature (red), and location: Anticosti (star), Florida Keys (triangle), North Mainland (circle), Saint Pierre et Miquelon (diamond) and South Mainland (square).

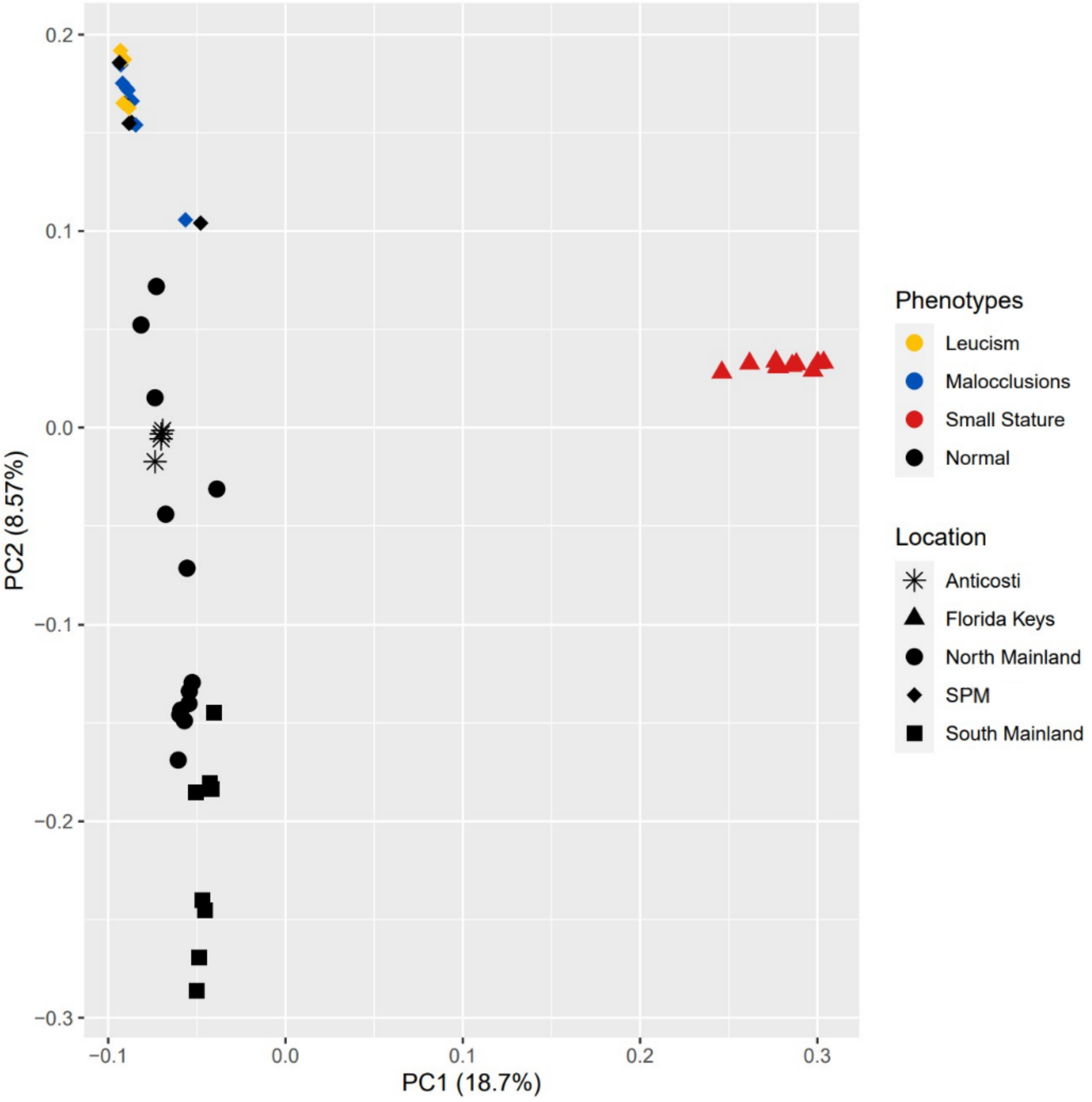

**Fig. S3:** High, Moderate, and Low influence homozygous mutations of each population based on SNPeff annotations. Populations are colour coded by phenotype: Normal (black), Leucism (yellow), Malocclusion (blue), and Small Stature (red).

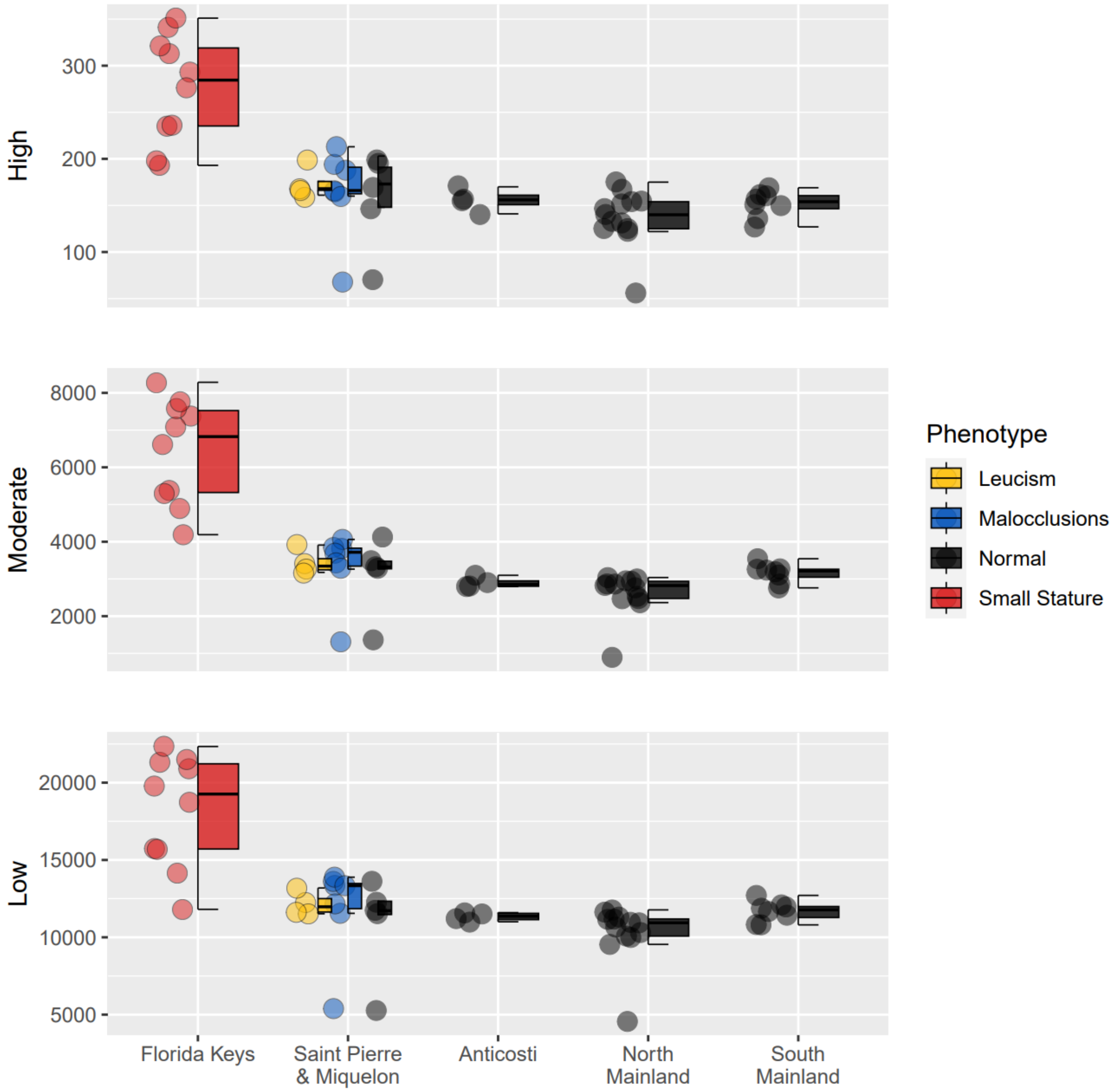

**Fig. S4:** Manhattan plot depicting results from the Genome-Wide Association Study (GWAS) conducted in PLINK between the leucism animals on St-Pierre & Miquelon and mainland populations. Genetic variants are shown along chromosomes (x-axis) by base pair (BP). Y-axis displays -log10 p-values, highlighting significant associations. Blue points denote specific fixed SNPs of interest and green points denote the BPs within the unique ROH pool of Leucism individuals.

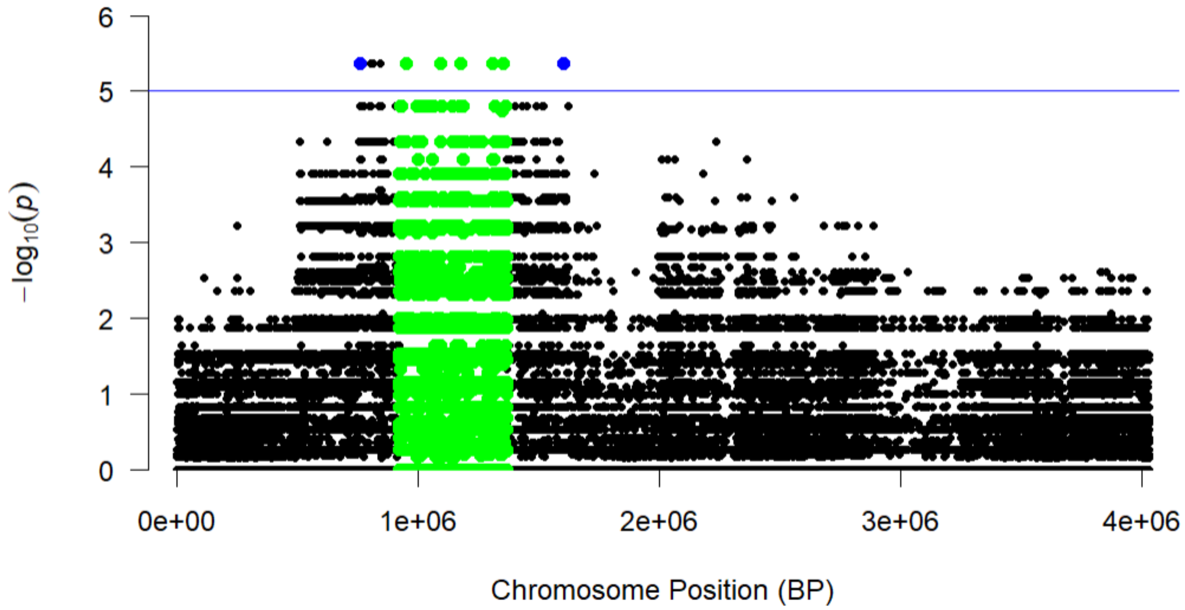

### Table S1 and Table S2 References


Bohn, G. *et al.* (2007) 'A novel human primary immunodeficiency syndrome caused by deficiency of the endosomal adaptor protein p14', *Nature Medicine*, 13(1), pp. 38–46. Available at: https://doi.org/10.1038/nm1528.

Chen, F. *et al.* (2015) 'Identification of a Mutation in FGF23 Involved in Mandibular Prognathism', *Scientific Reports*, 5, p. 11250. Available at: https://doi.org/10.1038/srep11250.

Guan, X. *et al.* (2015) 'The ADAMTS1 Gene Is Associated with Familial Mandibular Prognathism', *Journal of Dental Research*, 94(9), pp. 1196–1201. Available at: https://doi.org/10.1177/0022034515589957.

Haase, B. *et al.* (2013) 'Accumulating Mutations in Series of Haplotypes at the KIT and MITF Loci Are Major Determinants of White Markings in Franches-Montagnes Horses', 8(9), pp. e75071-e75071. Available at: https://doi.org/10.1371/journal.pone.0075071.

Hauswirth, R. *et al.* (2012) 'Mutations in MITF and PAX3 Cause "Splashed White" and Other White Spotting Phenotypes in Horses', *PLOS Genetics*, 8(4), p. e1002653. Available at: https://doi.org/10.1371/journal.pgen.1002653.

Hofstetter, S. *et al.* (2019) 'A non-coding regulatory variant in the 5′-region of the MITF gene is associated with white-spotted coat in Brown Swiss cattle', 50(1), pp. 27–32. Available at: https://doi.org/10.1111/age.12751.

Kaelin, C.B. *et al.* (2012) ‘Specifying and Sustaining Pigmentation Patterns in Domestic and Wild Cats’, *Science (New York, N.Y.)*, 337(6101), pp. 1536–1541. Available at: https://doi.org/10.1126/science.1220893.

Liu, S. *et al.* (2022) ‘A high-quality assembly reveals genomic characteristics, phylogenetic status, and causal genes for leucism plumage of Indian peafowl’, *GigaScience*, 11, p. giac018. Available at: https://doi.org/10.1093/gigascience/giac018.

Nikopensius, T. *et al.* (2013) ‘A Missense Mutation in DUSP6 is Associated with Class III Malocclusion’, *Journal of Dental Research*, 92(10), pp. 893–898. Available at: https://doi.org/10.1177/0022034513502790.

Perillo, L. *et al.* (2015) ‘Genetic Association of ARHGAP21 Gene Variant with Mandibular Prognathism’, *Journal of Dental Research*, 94(4), pp. 569–576. Available at: https://doi.org/10.1177/0022034515572190.

Reiner, G., Weber, T., *et al.* (2020) ‘A genome-wide scan study identifies a single nucleotide substitution in MC1R gene associated with white coat colour in fallow deer (Dama dama).’, 21(1). Available at: https://doi.org/10.1186/s12863-020-00950-3.

Reiner, G., Tramberend, K., *et al.* (2020) ‘A genome-wide scan study identifies a single nucleotide substitution in the tyrosinase gene associated with white coat colour in a red deer (Cervus elaphus) population’, *BMC Genetics*, 21. Available at: https://doi.org/10.1186/s12863-020-0814-0.

Rodrigues, J.B. *et al.* (2013) ‘Analysis of new Matrilin-1 gene variants in a case-control study related to dental malocclusions in Equus asinus’, *Gene*, 522(1), pp. 70–74. Available at: https://doi.org/10.1016/j.gene.2013.03.084.

Sun, R. *et al.* (2018) ‘Identification and Functional Studies of MYO1H for Mandibular Prognathism’, *Journal of Dental Research*, 97(13), pp. 1501–1509. Available at: https://doi.org/10.1177/0022034518784936.

Wang, C. *et al.* (2015) ‘Genome-wide analysis reveals artificial selection on coat colour and reproductive traits in Chinese domestic pigs’, *Molecular Ecology Resources*, 15(2), pp.

Wilderman A, D’haene E, Baetens M, Yankee TN, Winchester EW, Glidden N, et al. (2022). A distant global control region is essential for normal expression of anterior HOXA genes during mouse and human craniofacial development. bioRxiv: 2022.03.10.483852.414–424. Available at: https://doi.org/10.1111/1755-0998.12311.

Xu, L. *et al.* (2015) ‘Genomic Signatures Reveal New Evidences for Selection of Important Traits in Domestic Cattle’, *Molecular Biology and Evolution*, 32(3), pp. 711–725. Available at: https://doi.org/10.1093/molbev/msu333.

Xue, F., Rabie, A.B.M. and Luo, G. (2014) ‘Analysis of the association of COL2A1 and IGF-1 with mandibular prognathism in a Chinese population’, *Orthodontics & Craniofacial Research*, 17(3), pp. 144–149. Available at: https://doi.org/10.1111/ocr.12038.

Yu H, Wang Y, Gao J, Gao Y, Zhong C, Chen Y (2023). Application of the neuropeptide NPVF to enhance angiogenesis and osteogenesis in bone regeneration. Commun Biol 6: 1–12.
